# Chaos and the (un)predictability of evolution in a changing environment

**DOI:** 10.1101/222471

**Authors:** Artur Rego-Costa, Florence Débarre, Luis-Miguel Chevin

## Abstract

Among the factors that may reduce the predictability of evolution, chaos, characterized by a strong dependence on initial conditions, has received much less attention than randomness due to genetic drift or environmental stochasticity. It was recently shown that chaos in phenotypic evolution arises commonly under frequency-dependent selection caused by competitive interactions mediated by many traits. This result has been used to argue that chaos should often make evolutionary dynamics unpredictable. However, populations also evolve largely in response to external changing environments, and such environmental forcing is likely to influence the outcome of evolution in systems prone to chaos. We investigate how a changing environment causing oscillations of an optimal phenotype interacts with the internal dynamics of an eco-evolutionary system that would be chaotic in a constant environment. We show that strong environmental forcing can improve the predictability of evolution, by reducing the probability of chaos arising, and by dampening the magnitude of chaotic oscillations. In contrast, weak forcing can increase the probability of chaos, but it also causes evolutionary trajectories to track the environment more closely. Overall, our results indicate that, although chaos may occur in evolution, it does not necessarily undermine its predictability.

## Introduction

The extent to which evolution is repeatable and predictable bears on the usefulness of evolutionary biology as a tool for a growing number of applied fields, including drug resistance management in pests and pathogens, or sustainable agriculture and harvesting under climate change. So far, the investigation of factors that may reduce the predictability of evolution has mostly focused on various sources of stochasticity (i.e. randomness), namely genetic drift, the contingency of mutations, and randomly fluctuating environments (Crow & Kimura, 1970; Lenormand et al., 2009; Sæther & Engen, 2015). A much less explored source of unpredictability in evolution is deterministic chaos (but see Hamilton, 1980; Altenberg, 1991; Gavrilets & Hastings, 1995; Abrams & Matsuda, 1997; Dercole & Rinaldi, 2010; Dercole et al., 2010; Doebeli & Ispolatov, 2014), which occurs when the dynamics of a system, despite being completely predictable from the initial conditions, are strongly sensitive to them. Under chaotic dynamics, any measurement error, regardless how small, will be amplified over time, to the point that it becomes impossible to make accurate predictions beyond a certain timescale (Ott, 2002; Petchey *et al*., 2015).

It was recently demonstrated theoretically that evolutionary dynamics at the phenotypic level can become chaotic when natural selection results from between-individual interactions within a species (Doebeli & Ispolatov, 2014), i.e., even in the absence of any interspecific eco-evolutionary feedbacks, which are known to enhance ecological chaos (Abrams & Matsuda, 1997; Dercole & Rinaldi, 2010; Dercole *et al*., 2010). Specifically, evolutionary chaos arises in single-species models when (*i*) selection is frequency-dependent, such that the fitness of an individual depends on trait-mediated interactions with conspecifics; (*ii*) the fitness effects of these interactions are not simply a function of the phenotypic difference between the interactors (unlike typical models of within- and between-species interactions, e.g. Dieckmann & Doebeli, 1999; Doebeli & Ispolatov, 2010); and (*iii*) the number of traits involved in these interactions (described as organismal complexity) is large (Doebeli & Ispolatov, 2014).The authors concluded from this study that evolution is likely to be much less predictable than generally perceived. However, the theoretical demonstration that chaos is possible in a system is not sufficient to argue that this system is necessarily unpredictable, as we elaborate below.

First, the parameter values that lead to chaos may be rare in nature (Hastings et al., 1993; Zimmer, 1999). For instance in ecology, chaos has long been known to be a possible outcome of even simple population dynamic models (May, 1976), but despite a few clear empirical demonstrations in the laboratory (Benincá *et al*., 2008) and in the wild (Benincá et al., 2015), most natural populations seem to have demographic parameters placing them below the “edge of chaos”(Hastings *et al*., 1993; Ellner & Turchin, 1995; Zimmer, 1999; Dercole *et al*., 2006).

Second, and importantly with respect to evolution, a potentially chaotic system may still be predictable because it is subject to forcing by external factors with autonomous dynamics. In eco-evolutionary processes, such external forcing generally results from a changing environment that affects fitness components and their dependence on phenotypes, causing variation in natural selection. In fact, environmental variability affecting population growth (Lande *et al*., 2003; Ellner et al., 2011; Pelletier *et al*., 2012) and natural selection (MacColl, 2011; Chevin *et al*., 2015) is probably ubiquitous in natural populations, as documented notably by numerous examples of ecological and evolutionary responses to climate change (reviewed by Davis et al., 2005; Parmesan, 2006; Hoffmann & Sgrò, 2011). When such external forcing is imposed, the predictability of evolution is likely to change, because (*i*) forcing can alter the probability that the system is indeed chaotic, for instance by suppression of chaos through synchronization to the external forcing (e.g. Pikovsky *et al*., 2003); and, (*ii*) even if the dynamics remain chaotic, they may still be affected by the forcing in ways that make them largely predictable.

We investigate how a changing environment affecting phenotypic selection, modeled as a moving optimal phenotype (a classic approach, reviewed by Kopp & Matuszewski, 2014), influences the predictability of evolutionary dynamics in a context where chaos is expected to make evolution highly unpredictable in the absence of environmental forcing. Focusing on a periodic environment, we ask how the amplitude and period of cycles influence (*i*) the probability that evolutionary trajectories are chaotic, and (*ii*) the degree to which those trajectories that are indeed chaotic track the optimal phenotype set by the environment, making them partly predictable.We show that a changing environment can dramatically alter the probability of evolutionary chaos in either direction, but that evolutionary tracking of the environmental forcing generally contributes to making evolutionary trajectories much more predictable than anticipated from a theory that ignores any role of the external environment. This suggests that the predictability of evolution is partly determined by a balance between the strength of intraspecific interactions and responses to environmental change.

## Methods

### Model

We consider a set of *d* phenotypic traits that evolve under both frequency-dependent and frequency-independent selection. Frequency-independent selection is assumed to be caused by stabilizing selection towards an optimal trait value, at which carrying capacity *K* is maximized. For instance, selection on beak size/shape in a bird, or mouth shape in a fish, may have a local optimum set by the available type of resources (Martin & Wainwright, 2013; Grant & Grant, 2014). The frequency-dependent component of selection, on the other hand, emerges from trait-mediated ecological and social interactions between individuals within the species (either cooperative or competitive). Selection on a bird’s beak morphology, for instance, depends not only on the available types of resources, but also on competition with conspecifics for these resources (as in Grant & Grant, 2014). The intensity of this competition may depend on the beak size of competing individuals, but also on other traits of these interactors, such as their aggressiveness, territoriality, or degree of choosiness in food preference. This frequency-dependent component of selection, when it involves many traits, can lead to complex dynamics such as chaos or internally driven cycles, as shown by Doebeli & Ispolatov (2014).

Other details of the models, laid out in the Supporting Information, follow Doebeli & Ispolatov (2014) for ease of comparison. Notably, we use the same adaptive dynamics assumptions, whereby evolution is slower than population dynamics and relies on rare new mutations (Dieckmann & Law, 1996; Geritz *et al*., 1998; Dercole & Rinaldi, 2008), although our results are likely to apply also in a quantitative genetic context where evolutionary dynamics are much faster (see Discussion). Under these assumptions, the evolutionary dynamics of each phenotypic trait *X_i_* is (see Supporting Information for more details)

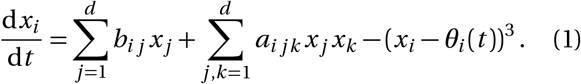

The first two terms in equation (1) represent frequency-dependent selection caused by phenotype-dependent interactions between individuals. The coefficients *b_ij_* and *a_ijk_* determine, respectively, the strength of first- and second-order selective interactions between traits. The latter occurs when the fitness of an individual with a given phenotype at trait *i* depends on the product of its phenotypic values at two traits *j* and *k* (including *j* = *i* and/or *k* = *i*), at least one of which is from its interactor (e.g. beak size of focal individual interacting with beak size and agressiveness of interactor). Importantly, these frequency-dependent components of directional selection will be null if the interaction between individuals depends only on their phenotypic difference, in which case frequency-dependent selection would not lead to chaotic dynamics (as explained in the Supporting Information). Put differently, this means that a necessary (but not sufficient) condition for evolutionary chaos in this model is that intraspecific interactions do not depend solely on the resemblance (or difference) between the trait values of interactors, but also on their actual phenotype, for instance when individual that are highly social, agressive, or large, interact more overall.

The last term in equation (1) models stabilizing selection that causes the phenotype to evolve towards the optimum *θ_i_* for each trait *i*. In the original model (Doebeli & Ispolatov, 2014), the optimal phenotype was assumed to be constant and equal to zero for all traits. Here in contrast, we modeled the forcing effect of a changing environment by letting the optimum ***θ*** (*t*) change with time. We focused on a cycling environment causing the optimum to oscillate sinusoidally, which may represent, depending on the organism, oscillations in biotic (predators, parasites) or abiotic (e.g. meteorological) conditions on seasonal, multi-annual (e.g., El Niño oscillation), or geological time scales. We assume for simplicity that the optimal values for all traits respond to the same underlying environmental variable, such that they oscillate with the same period and phase (i.e., they are synchronized). The vector of optimal phenotypes for all traits can then be written as

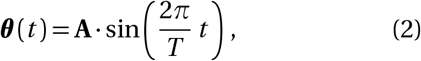

where *T* is the period, and **A** = (*A*_1_, *A*_2_, …, *A_d_*) is a vector of amplitudes of oscillation for each of the *d* traits. This defines a single sine wave of amplitude ║**A**║ (the norm of vector A), and direction given by the unitary vector **A**/║**A**║. In our simulations, we focused for simplicity on the case where *A_i_* is the same for all traits, such that the optimum changes along a diagonal of the phenotype space. Note also that equation (2) implies that the optimum fluctuates around the phenotype for which the strength of selective interactions vanishes (set to the origin by definition, without loss of generality). Allowing for fluctuations to be centered on a different phenotype – or equivalently, including a 0^*th*^ order term in the interaction component of selection in equation (1) – would select for increasing interactions in all environments, thus artificially increasing the probability of chaos relative to Doebeli & Ispolatov (2014), where the optimum was set constant at the origin.

### Simulations

We studied the evolution of the phenotypic vector **x**(*t*) containing the mean phenotype for each trait by numerically solving the dynamics of equation (1) given the initial phenotype **x**(0), period *T* and vector of amplitudes **A** (both characterizing the optimum ***θ***), and set of interaction coefficients *a_ijk_* and *b_ij_*. In each simulation, the interaction coefficients were independently drawn from a normal distribution with mean 0 and standard deviation 1, and rescaled as 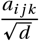 and 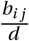. This rescaling, proposed by Doebeli & Ispolatov (2014), ensures that dynamics under different dimensionalities under a constant environment explore similar ranges of phenotypic values, between—1 and 1. It also prevents the unrealistically strong selection produced at high dimensionalities in the model with unscaled interaction coefficients: in effect, we keep the expected overall strength of selection constant but spread it across the *d* traits. Trajectories were run up to *t* = 1200 using the LSODA method, as implemented in the package deSolve in R (Soetaert *et al*., 2010; R Core Team, 2015), with integration step dt = 0.1. Initial phenotypes were drawn from a multivariate normal distribution centered at zero such that, on average, the carrying capacity *K*(_x_(0), *_θ_*(0)) = 0.5 (Eq. S2). This choice was made to keep the initial state of the system under biologically relevant degrees of adaptation.

Evolutionary dynamics were categorized based on their largest Lyapunov exponent λ, which measures the rate of exponential increase in the distance between two trajectories that start from very close initial conditions (Sprott, 2001). Dynamics that converge to an equilibrium phenotype have *λ* < 0, those that oscillate periodically (limit cycles) have *λ* = 0, and those that systematically diverge due to strong sensitivity to initial conditions (which defines chaos) have *λ* > 0. Here we used a local average Lyapunov exponent computed over a sliding window of 200 time units (see Supporting Information and Supplementary Fig.S1).

### Proportion of transient chaos in a constant environment

In this model, trajectories that eventually reach fixed points or limit cycles may exhibit complex behaviors for long periods of time (Fig.1A and B), during which they are indistinguishable from chaos. To understand the prevalence in the system of this so-called *transient chaos*(Grebogi *et al*., 1983; Gavrilets & Hastings, 1995; Lai & Tél, 2010), we ran simulations with *d* ranging from 2 to 100 traits, under a constant optimum set at zero for all traits. Trajectories were identified as transiently chaotic if they switched from *λ* > 0 to either *λ* = 0 or *λ* < 0 (i.e. they went from chaos to a cycle or fixed equilibrium, respectively) by the end of the simulations (at time *t* = 1200; excluding the first 50 time steps of the simulations that correspond to a burn-in period).

**Fig. 1.**
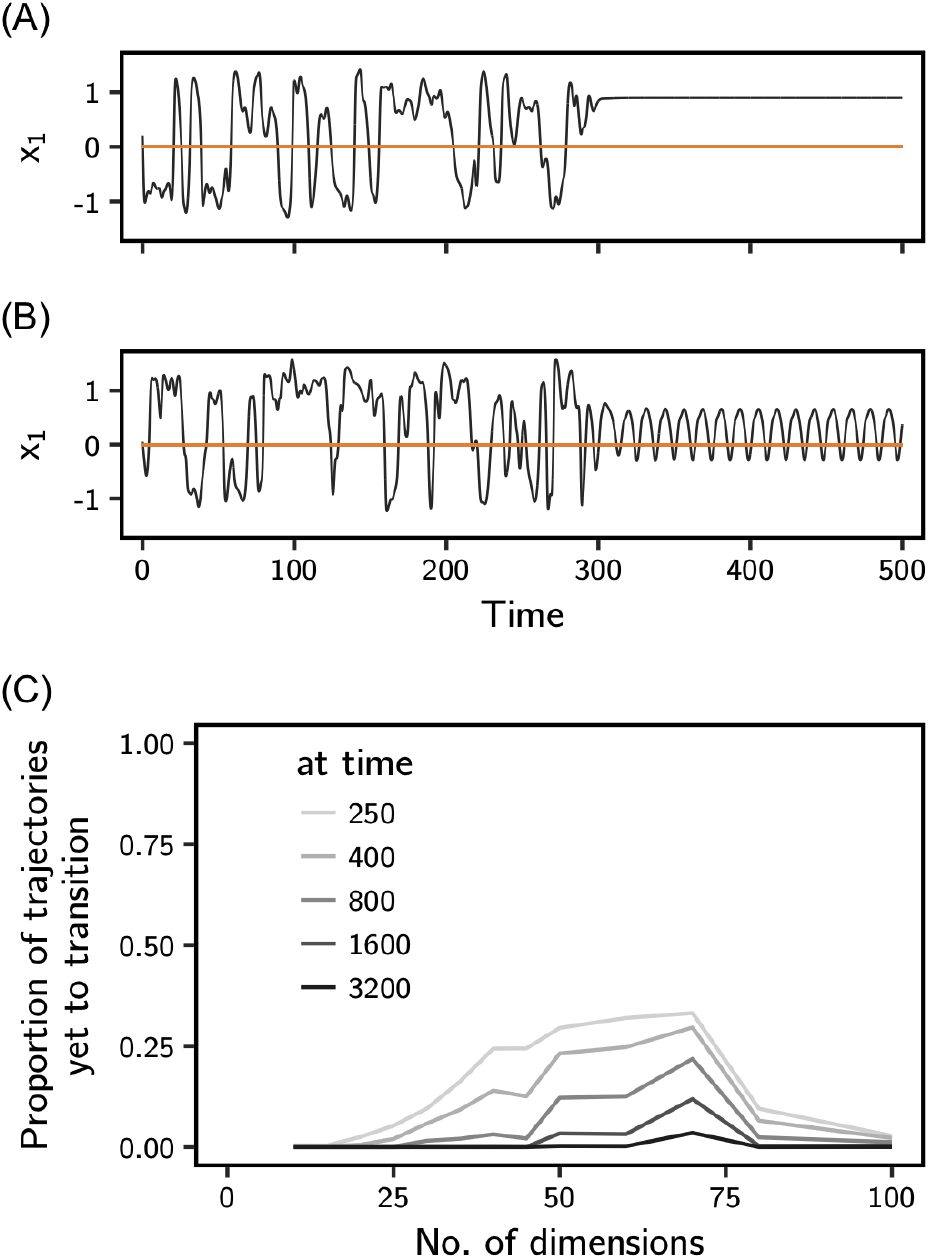
Transients in a constant environment. Transient trajectories present chaos-like behavior for some time, before transitioning to either (A) fixed equilibrium phenotypes or (B) periodic cycles. Two representative trajectories for a single trait are shown, simulated as described in the Methods, with *d* = 45 and a constant optimum (orange line). (C) The expected proportion of evolutionary trajectories (among all those for a given dimensionality *d*) that will eventually transition to non-chaotic dynamics, but would still be categorized as chaotic at a given time, are shown for different times. These proportions were estimated from a statistical model of exponential decrease with time of the proportion of apparently chaotic dynamics (model fits in Supplementary Fig.S2). For each dimensionality *d*, 250 trajectories were run up to *t* = 1200 and classified based on their estimated Lyapunov exponents λ, as described in the Methods.

Transience is expected to cause the proportion of chaos-like trajectories to decrease exponentially with time (Yorke & Yorke, 1979), even for high-dimensional systems (Grebogi *et al*., 1983). For each dimensionality, we thus estimated the proportion *f*(*t*) of trajectories behaving chaotically at each time step, and used this to estimate the asymptotic proportion of trajectories that remain chaotic over infinite time. This was done using the non-linear least-squares method (`nls` function in R’s *stats* package R Core Team, 2015) to fit a statistical model of the form *f*(*t*) = *A* exp(*B t*)+*C*+*ϵ*, where *A*, *B* and *C* are the estimated variables (*C* being the asymptotic proportion of chaos), and *ϵ* is the residue (see Supplementary Fig.S2).

### Predictability in a changing environment

To investigate the effect of a changing environment, we focused on a system of high dimensionality (*d* = 70), because this leads to a high probability of chaos in a constant environment (Supplementary Fig.S3 and Doebeli & Ispolatov, 2014). We used 100 sets of interaction coefficients *b_ij_* and *a_ijk_* and initial phenotypes **x**_0_. Each set of parameters was used for simulations in a constant environment and in changing environments (40 different combinations of amplitude ║**A**║ and period *T* of optimum oscillation, with ║**A**║ ∈ {1.30, 2.33, 3.73, 5.04, 5.99} and *T* ∈ {1.5, 2, 3, 5, 10, 20, 50, 100}.) The amplitudes were chosen such that the smallest carrying capacity that a phenotype centered at the origin would experience (i.e. when the optimal phenotype was at the peak of its oscillation) was {0.99, 0.9, 0.5, 0.1, 0.01}.

For each regime of environmental change, we investigated the extent to which chaotic trajectories (i.e. trajectories with a final Lyapunov exponent *λ* > 0) track the moving optimum. For this, the time-series of phenotypic values were regressed on the oscillating optimal phenotype. We focused on the projection 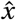 of the multivariate phenotypes along the direction of oscillations of the optimum in the phenotypic space, which we regressed on a similar projection 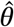 for the optimum (which is simply the norm of ***θ***). Additionally, because the evolving phenotype systematically lags behind the moving optimum in such a system (Lande & Shannon, 1996), we maximized the *R*^2^ of the regression by shifting forward the time-series of phenotype relative to that of the optimum (as shown in Supplementary Fig.S4). This maximum *R*^2^ quantifies the proportion of variance in the evolutionary dynamics explained by (lagged) tracking of the optimum, so it is a measure of the predictability of evolution conditional on knowledge of the environment. This tracking component of evolution necessarily has the same period as the optimum (Supplementary Fig.S4), so approximating it as a sinusoidal function for simplicity, the regression slope of the phenotype 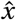 on the (resynchronized) optimum 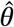 is just the ratio of amplitudes of their cycles (as we confirmed in our simulations, see Supplementary Fig.S5).

We also looked for signatures of tracking of the changing environment in the evolutionary time-series 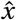 using spectral analysis (fast Fourier transform method, as implemented in R’s *stats* package, R CoreTeam, 2015). This technique treats time-series as superpositions of sine and cosine waves of different frequencies (Shumway & Stoffer, 2011), and allows estimation of the amplitudes associated to each frequency of oscillation (1/*T*). More precisely, the spectral density as computed by the `spectrum()` function in R is half the squared amplitude of waves of the corresponding frequency of oscillation.

## Results

### Transient evolutionary chaos is common

Chaos in phenotypic evolution was previously shown to be a common outcome in a constant environment, when interactions with conspecifics cause frequency-dependent selection as modeled in equation (1) and the interactions are mediated by many traits (Doebeli & Ispolatov, 2014; Ispolatov *et al*., 2015). However, some of the apparently chaotic trajectories are actually transient (Fig.1A and B). The fraction of trajectories that exhibit such transient chaos strongly depends on organismal complexity *d* (Fig.1C), being highest for *d* between 40 and 70 (approximately), where the proportion of trajectories that are chaotic is intermediate (Fig. S2, and Doebeli & Ispolatov, 2014; Ispolatov *et al*., 2015). The statistical models fitted to the proportion of chaotic trajectories in time predict that virtually all transient trajectories should eventually transition to fixed points or periodic cycles by *t* = 3200 (Fig.1C). However before this transition, the phenotype may undergo very complex, random-like dynamics, with no indication that they will eventually transition to a state that, once established, is quite predictable (see Fig.S3).

### A changing environment may either increase or decrease the probability of chaos

We next investigated the effect of a changing environment. We tracked the proportion of trajectories that were chaotic (up to time *t* = 1200) under sinusoidal cycles in the optimal phenotype, with varying periods and amplitudes. The proportion of chaotic trajectories was strongly influenced by the regime of optimum oscillation (Fig.2A). Long periods of oscillation increased the probability of chaos relative to a constant environment, for the same organismal complexity (*d* = 70). For long enough periods, essentially all trajectories became chaotic at dimensionality *d* = 70, so the chaos-enhancing effect depended little on the amplitude of oscillations. However at smaller dimensionality (d = 40), the probability of chaos was maximized for a period that depended on the amplitude of oscillations, decreasing for very large periods when the amplitude was large (Fig.S6).

**Fig. 2.**
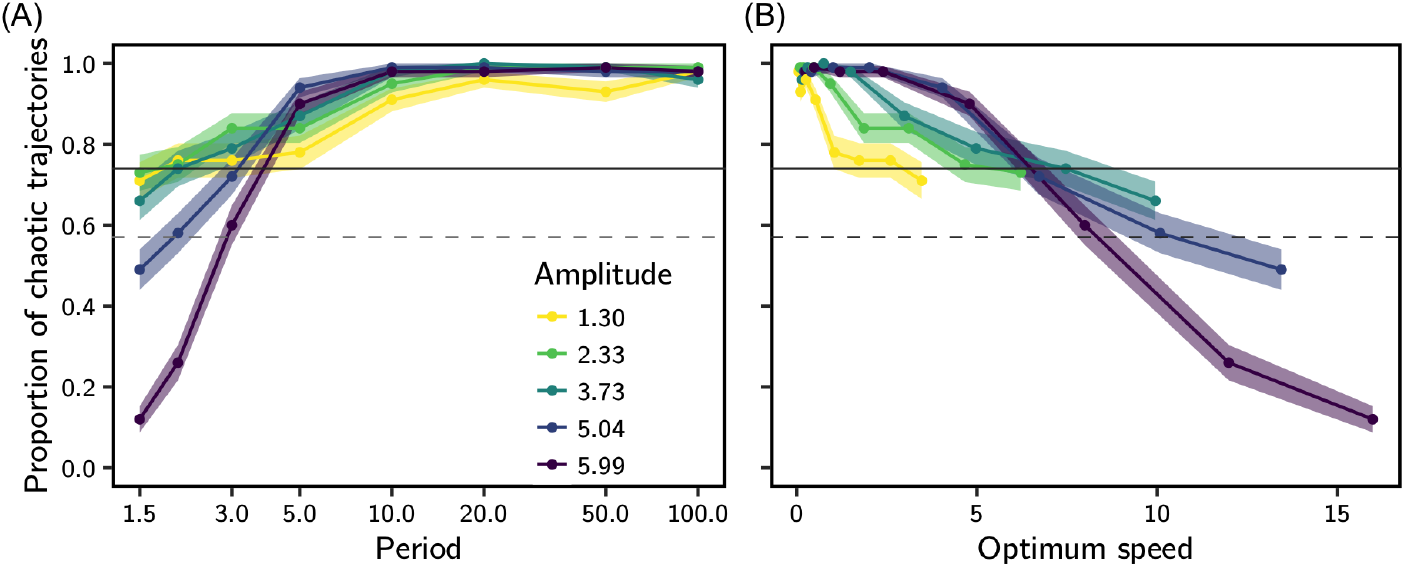
Probability of chaos in a changing environment. The proportion of chaotic trajectories at time *t* = 1200 in an oscillating environment is shown against (A) the period and (B) the average speed of optimum oscillation, for different values of amplitude (colors). For each condition of optimum oscillation (period and amplitude), we report the estimate (dot, line) and standard error (shading) of the proportion of chaotic trajectories, out of 100 replicated simulations that used sets of parameters also used in the constant-environment simulations (with dimensionality *d =* 70). The average speed of optimum oscillation is calculated from equation (2) as 4║**A**║/*T*. The observed proportion of chaos at *t* = 1200 (horizontal solid line), and the predicted proportion after all chaotic transients have transitioned (dashed line; calculated as shown in Supplementary Fig.S2) are also represented for a constant environment with *d* = 70. While long periods cause the proportion of chaotic trajectories to increase relative to a constant environment, short periods of large amplitudes (fast oscillations) cause this proportion to decrease, even below what would be predicted through more rapid transition of transiently chaotic trajectories.

In contrast to this chaos-enhancing effect, a combination of large amplitudes and short periods of optimum oscillation led to a sharp decrease in the proportion of chaotic trajectories (Fig.2A). This decrease at time *t* = 1200 is not simply caused by an earlier transition out of chaos by transient trajectories: the observed proportion of chaotic trajectories can be substantially lower than that projected in infinite time for a constant optimum condition.

Part of the effect of the changing environment on the probability of chaos is explained by the absolute speed of the moving optimum. A simple metric of speed is the norm of the derivative of equation (2) relative to time, whose average over a period is equal to 4║**A**║/*T*. We use this average value as our measure of optimum speed. A slow optimum (speed below 5 for *d* = 70, Fig.2B) increases the probability of chaos relative to a constant environment, while a fast optimum (speed above 5 for *d* = 70, Fig.2B) decreases the probability of chaos. Furthermore, conditions that lead to a reduced probability of chaos relative to a constant environment (frequent oscillations with broad amplitude) also correspond to strong environmental forcing on the evolutionary dynamics, causing substantial maladaptation and directional selection even in a context without frequency dependence (Lande & Shannon, 1996), as shown in Supplementary Fig.S7.

### Strong environmental forcing renders chaotic trajectories more predictable

The predictability of evolution is not entirely captured by the probability of chaos, even in a deterministic system as modeled here, because chaotic evolutionary trajectories need not be entirely unpredictable. In fact these trajectories, despite looking erratic, are constrained to remain near the optimal phenotype imposed by stabilizing selection (Doebeli & Ispolatov, 2014). When a changing external environment causes movements of the optimum, evolution necessarily tracks this moving optimum to some extent, even for chaotic trajectories, as illustrated in Fig.3A and B. The predictability of evolution for those trajectories that are chaotic thus depends on how the internally driven dynamics due to intraspecific interactions interplay with forcing by the external environment.

**Fig. 3.**
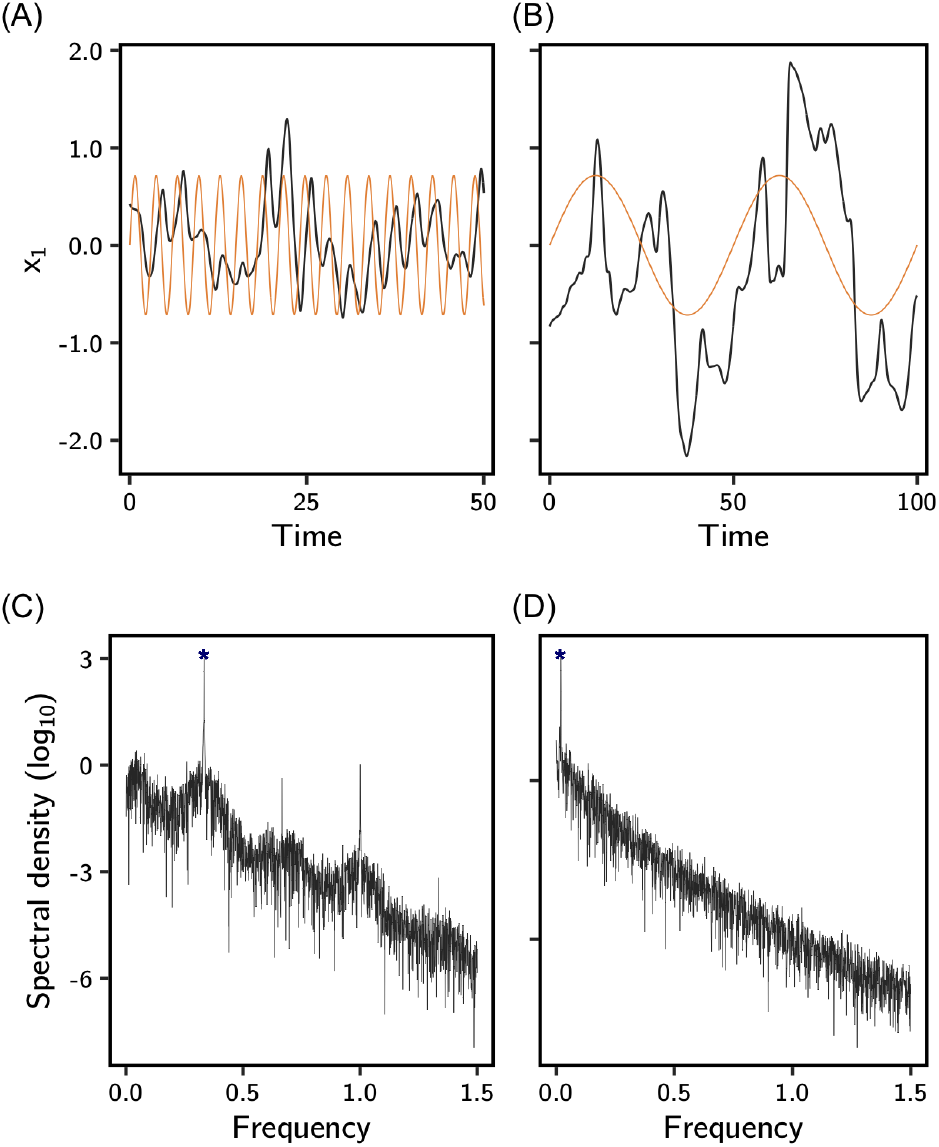
Chaotic evolutionary dynamics in changing environments. Chaotic evolutionary trajectories (black in A and B) combine internally driven chaos with external environmental forcing through tracking of the optimum. Chaotic oscillations can be either (A) longer or (B) shorter than the oscillations of the optimum (shown in orange), depending on the periods of the latter (3 and 50, respectively). (C, D) Spectral analysis of these same time-series of evolving phenotype (using all traits as described in the Methods) show a peak of spectral density (blue star) at the frequency of oscillation that corresponds to the oscillating optimum.

To investigate how tracking of the environment affects the predictability of evolution, we regressed the time series of the evolving phenotype on that of the optimum, for trajectories that are chaotic in a changing environment. Such regressions are similar to phenotype-environment associations, as are commonly estimated empirically from time series of phenotypes and environments (review in e.g. Michel *et al*., 2014), and similarly across space (e.g. Phillimore *et al*., 2010). Note that here, the phenotype first needs to be resynchronized with the environment, to correct for the adaptational lag (Fig.S4, Lande & Shannon, 1996). The *R*^2^ of the regression of phenotype on the environment is a measure of the predictability of evolutionary trajectories conditional on knowledge of the environment, since it measures the proportion of the total temporal variance in phenotype that is explainable by tracking of the optimum. The predictability of chaotic evolutionary dynamics increases with larger amplitudes and longer periods of optimum oscillation (Fig.4A; and see Supplementary Fig.S8 for regressions on optimum speed). Indeed, (*i*) evolutionary trajectories track long-period oscillating optima more closely than they do short-period ones (Lande & Shannon, 1996), and (*ii*) if this optimum undergoes ample fluctuations, so will the phenotype, such that the tracking component of evolution will be substantial, as illustrated in Fig.3B.

**Fig. 4.**
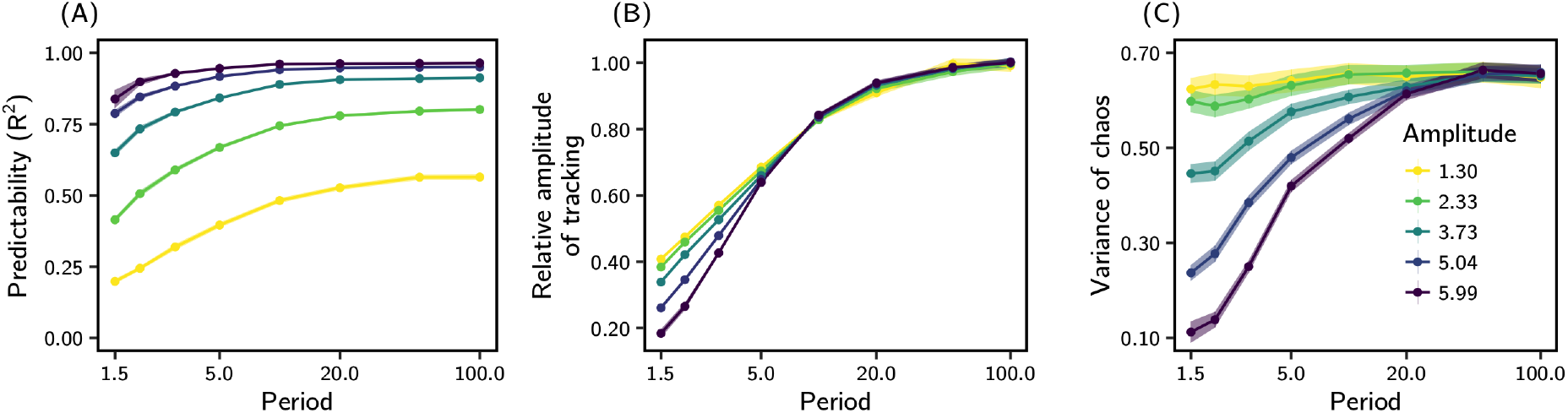
The linear regression of phenotype on the moving optimum gives insights about the predictability of chaotic evolutionary dynamics. (A) The fraction of the total temporal variation in the evolving phenotype explained by movements of the optimum, as captured by the *R*^2^ of the regression, is higher when the optimum oscillates with larger amplitudes and longer periods. (B) The ratio of amplitudes between the tracking component of oscillations in the evolving phenotype and oscillations in the optimum, as estimated by slope of the regression, is smaller under short-period optimum oscillations. (C) Temporal variation in evolutionary dynamics introduced by chaos, beyond the variation attributable to tracking of the moving optimum, is captured by the variance of residuals in the regression. Conditions that allow for close tracking of the optimum in (B) also lead to dampened chaotic oscillations. We show the average (lines and points) and standard error (shading) over 100 simulations for each combination of amplitude and period of optimum oscillation.

The exact same pattern is found for the repeatability of evolutionary trajectories among replicates, as measured by the proportion of the total variance in evolutionary trajectories attributable to temporal variation in the mean trajectory across replicates (Supplementary Fig.S9). The rationale for this measure is that the mean trajectory captures the tracking component of evolutionary change, while any additional variance around this trajectory is contributed by the internal dynamics caused by frequency-dependent selection. Repeatability thus has a very similar meaning to predictability, but unlike the latter it does not require information about the environment. Note also that repeatability is a measure of cross-predictability of evolution across replicates, and hence a metric of parallel phenotypic evolution, as commonly reported in the laboratory and in the field (Lenormand *et al*., 2016).

The amplitude of the tracking component of phenotypic evolution (scaled to the amplitude of the optimum oscillation) is quantified by the regression slope of phenotype on optimum. This relative amplitude increases with the period of cycles in the optimum, tending towards 1 for long-period cycles of any amplitude (Fig.4B). In contrast for shorter periods of optimum oscillation (left half of Fig.4B), evolutionary tracking of the environment is more moderate, and less efficient for larger amplitudes.

The chaotic, non-predictable component of the evolutionary dynamics is captured by the residuals of the regression of the phenotype on the optimum. The variance of residuals was lower under combinations of large amplitudes and small periods of environmental change (Fig.4C), corresponding to fast optima. Therefore, strong environmental forcing further increases the predictability of chaotic trajectories by dampening the extent to which they oscillate chaotically around the optimum. This effect can be seen in Fig.3A, where the trajectory departs less from the optimum (due to chaos) than the one in Fig.3B, with same amplitude but larger period.

Finally, we used spectral analysis, a general signal processing approach, to investigate whether environmental tracking leaves a signature in individual evolutionary time series. Similarly to the repeatability analysis, this involves the investigation of the evolutionary dynamics without knowledge of the forcing environment. For essentially all regimes of environmental change that we investigated, the frequency of oscillation with highest spectral density in the time series of phenotypic evolution corresponded to that of the oscillating optimum, as illustrated for specific cases in Fig.3C and D. The amplitude of the wave corresponding to this frequency of oscillation, as estimated from the spectral density, matched the amplitude estimated from the regression analysis (Fig.4B) very well (Supplementary Fig.S5). This shows that even in a context where evolutionary dynamics can be largely chaotic, the strongest signal in the evolutionary time series is likely to be the predictable response to environmental change.

## Discussion

When investigating factors that reduce predictability of evolution, evolutionary biologists have mostly focused on different sources of stochasticity (Crow & Kimura, 1970; Lenormand *et al*., 2009; Sæther & Engen, 2015), while the possibility of chaos has received comparatively less attention. Doebeli & Ispolatov (2014) have recently shown that interactions mediated by many traits can produce chaos in phenotypic evolution. However, the implications for the predictability of evolution should not be overemphasized. First, the condition for chaos to occur is not simply frequency-dependent selection on many traits: it also requires that the strength of intraspecific interactions depends directly on the phenotypes of interactors, rather than only on their phenotypic match or distance (Doebeli & Ispolatov, 2014), while interaction of the latter type are more commonly used in models of evolutionary ecology. Second, we reveal that chaos can be transient in this model, such that initially chaotic dynamics might not be observed after some time. And third, forcing by a changing environment, a ubiquitous driver of eco-evolutionary dynamics (as exemplified by responses to climate change, Davis *et al*., 2005; Parmesan, 2006; Hoffmann & Sgrò, 2011), can modify the predictability of evolution in a system that may be chaotic in a constant environment. Below we discuss this latter point further.

### Chaos vs. forcing

Strong environmental forcing (represented by a fast moving optimum) caused suppression of chaos relative to a constant environment in our model. In the general literature on chaos (outside of evolutionary biology), it has previously been demonstrated, both theoretically and experimentally, that chaotic dynamics can be suppressed with even slight forcing, but this usually occurs when the period of the forcing oscillation aligns closely to one of the specific resonance periods of the dynamics (Lima & Pettini, 1990; Fronzoni *et al*., 1991). In contrast in our simulations, the proportion of chaotic trajectories was reduced over a broad range of short periods and large amplitudes of oscillation that jointly result in a fast moving optimum. In fact, most of the chaos suppression occurred for periods much shorter than the typical internal chaotic dynamics in a constant environment (Supplementary Fig.S10). One of the reasons for this discrepancy may be that we studied the effect of the moving optimum on a total of 250 different sets of interaction coefficients, instead of focusing on single defined chaotic system, as usually done in the physics literature. Another reason is that in our evolutionary model, the dynamical system is different (and in general more complex) than in models studied in physics, for instance as in Lima & Pettini (1990) and Fronzoni *et al*.(1991).

In the opposite regime of slow optimum, the proportion of chaotic dynamics increased. Such an outcome is not uncommon in the general literature on chaos, as exemplified by the chaotic dynamics induced on a pendulum with friction by externally imposed sinusoidal impulses (forced damped pendulum, Ott, 2002). Such a phenomenon has also been shown in models of population ecology. Rinaldi et al.(1993) explored seasonal oscillation of demographic parameters in a predator-prey model, and found that such forcing easily led to quasiperiodic and chaotic dynamics that did not occur in a constant environment. Similarly, Benincá *et al*. (2015) designed a model based on an empirical rocky shore ecosystem with non-transitive interactions of the rock-paper-scissor form, and showed that the model only led to the chaotic oscillations observed in nature when including the forcing effect of yearly seasonal temperature cycles on death rates. In our model, optima that move slowly cause repeated long excursions away from trait values where interactions are minimal (**x** = 0 in the model). This causes stronger interaction terms in equation (1) (and hence more chaos) relative to a constant optimum at **x** = 0, because chaos-enhancing interactions are stronger with larger trait values in this model (Doebeli & Ispolatov, 2014).

### Generality and relevance to natural populations

Our results are difficult to compare quantitatively to empirical measurements, because we relied on the adaptive dynamics approach (to facilitate comparison with the original model in a constant environment by Doebeli & Ispolatov, 2014), where time is not measured in units of time or in generations, but in terms of an implicit number of mutations fixed. In principle, this would suggest that the timescales of evolutionary and environmental change in our model are restricted to be slow, since they are limited by the time between independent fixation events. However, several authors (Abrams *et al*., 1993; Waxman & Gavrilets, 2005; Débarre *et al*., 2014) have highlighted that the canonical equation of adaptive dynamics (on which eq. (1) is based) is almost identical to the equation for the response to selection in quantitative genetics (Lande, 1976), but with the latter operating on much shorter time scale. Furthermore, Ispolatov, Madhok & Doebeli (2016) demonstrated that the chaotic attractors of the original Doebeli & Ispolatov (2014) paper can be closely matched by individual-based simulations that allow for substantial polymorphism, and thus for much faster evolutionary dynamics than under the classic adaptive dynamics assumptions (*i.e*., rare substitution events in otherwise monomorphic populations). This suggests that the relevant timescale of evolutionary and ecological change in our model can be much faster than assumed by adaptive dynamics, e.g. that of microbial evolutionary experiments or long-term field surveys, for which it is well known that evolutionary change can happen quickly (e.g.Hendry & Kinnison, 1999; Campbell-Staton *et al*., 2017). Finally, understanding of the timescale of these chaotic oscillations will help elucidate the importance of transience, which should depend on the probability that an evolving transient population is observed before it has time to transition.

A crucial parameter of the model that is difficult to measure empirically is organismal complexity. The number of traits that can be measured is virtually infinite, but some may be highly correlated, or have negligible effects on fitness. From an evolutionary perspective, phenotypic complexity thus has to be defined with respect to its effects on fitness and selection. Previous theory has shown that complexity defined in a similar way as here (number of traits under stabilizing selection) has important impacts on the rate of adaptation (Fisher, 1930; Orr, 2000), speciation and diversification (Doebeli & Ispolatov, 2010; Chevin *et al*., 2014; Débarre *et al*., 2014; Svardal *et al*., 2014), or the drift load in a finite population (Poon & Otto, 2000; Tenaillon *et al*., 2007). Some of these predictions have been used to attempt to estimate organismal complexity indirectly, through its emerging effects on fitness effects of mutations. Very different results have been obtained, with complexity ranging from very low (on the order of 1) to several orders of magnitude for the same organism, depending of the underlying model used (Martin & Lenormand, 2006; Tenaillon *et al*., 2007). In any case, our results under a changing environment should apply whenever the complexity of frequency-dependent selection caused by intraspecific interactions is high enough to generate chaotic dynamics in a constant environment (Doebeli & Ispolatov, 2014).

A possibility not explored in our study is that a changing environment alters the interaction strength between individuals with given phenotypes, and thus the frequency-dependent component of selection in equation (1). It is not entirely clear how to best model such an effect of the environment on interaction coefficients – while a moving optimum for the frequency-independent component of selection is well-established in models (reviewed by Kopp & Matuszewski, 2014), and has some empirical support (Chevin *et al*., 2015). We can however anticipate that environments leading to larger absolute values of the interaction coefficients should increase the probability of chaos.

### Interacting sources of unpredictability in evolution

We have focused for simplicity on effectively infinite populations in a fully deterministic environment, such that any unpredictability in evolution has to come from sensitivity to initial conditions characteristic of chaotic dynamics. This is perhaps a good approximation for some experimental studies with microbes in controlled environments, but more generally populations in the wild should also be exposed to another source of unpredictability: evolutionary stochasticity caused by genetic drift, the contingency of mutations, or a randomly changing environment (Crow & Kimura, 1970; Lenormand *et al*., 2009; Sæther & Engen, 2015). A more complete understanding of the predictability of evolution would therefore require combining stochasticity and chaos to investigate their possible interactions, as advocated previously for population dynamics (Ellner & Turchin, 1995). For instance, stochastic factors have been shown to either increase or decrease (depending on the chaotic system) the time that transients take to converge to their equilibrium states (Lai & Tél, 2010).

While such an analysis is beyond our scope here, some preliminary statements can be made based on our results and those from the literature. Beyond just adding random variation among replicates (and thus directly reducing evolutionary predictability), stochasticity may interact with chaos in evolutionary dynamics, amplifying or reducing its importance. A stationary stochastic environment is a type of forcing that shares some similarities with deterministic cycles. Indeed, quantitative genetic models (Lynch & Lande, 1993; Lande & Shannon, 1996) have shown that increasing the stationary variance (respectively autocorrelation) of a stochastic environment has similar effects on the lag load (caused by phenotypic mismatches with the optimum) as increasing the amplitude (respectively period) of cycling environment. The results we report here could thus be used to guide interpretation about the outcomes from future models investigating the combined effects of stochasticity and chaos on the predictability of evolution. An interesting challenge for empirical research would be to establish the time scales at which chaos versus stochasticity dominate as sources of unpredictability in evolution.

## Acknowledgements

We thank R. Gomulkiewicz, S. de Monte, and J. Pantel for insightful discussions, and three anonymous reviewers for useful feedback on a previous version of the manuscript. ARC is funded by an Erasmus Mundus Joint Master Degree scholarship from the European Comission. FD is funded by a grant from Agence Nationale de la Recherche (ANR-14-ACHN-0003-01). LMC is funded by a grant from the European Research Council (ERC-2015-STG-678140-FluctEvol).

## Data Archiving

A minimum code for simulation and data analysis is currently available at https://figshare.com/s/6637d1e0c6eb320e3247 and will be uploaded to Dryad.

## Supplementary Information

### Details of the model

We extend the evolutionary model in a constant environment by Doebeli & Ispolatov (2014) to a fluctuating environment. The model takes as a starting point logistic population growth where the strength of density dependence depends on phenotype-mediated interactions,

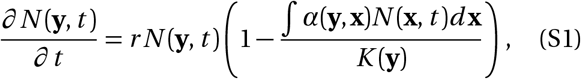

where *N*(**y**, *t*) is the number of individuals of type **y** at time *t*. The phenotype of an individual is represented by a vector **x** (or **y**) of length *d*, the number of traits under selection; *d* is called the dimensionality or complexity of the organism (Doebeli & Ispolatov, 2014). Interactions are mediated by phenotypes: an interaction kernel α(**y**, **x**) represents the fitness effects of interactions with individuals of phenotype **x** on individuals of phenotype **y**. These interactions can either be positive (e.g. cooperation) or negative (e.g. competition) depending on the sign of α (negative and positive, respectively). The interaction kernel is standardized such that α(**y,y**) = 1 for all **y**, but the exact form of the interaction kernel does not have to be specified at the moment. Another component of fitness is due to adaptation to the current environmental condition. It takes the form of a phenotype-dependent carrying capacity *K*, causing stabilizing selection towards a multivariate optimal phenotype *θ*. Specifically, we have

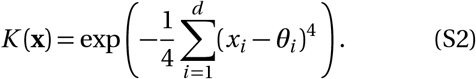

We use a quartic function because it is the form assumed by Doebeli & Ispolatov (2014). A more general form could include weights for the different traits, and even interactions between the traits, but we want to stay close to their original model.

The size of the entire population is assumed to be large; reproduction is clonal; mutations are rare, and of small phenotypic effect. To study the evolution of the mean phenotype in the population, and to facilitate comparison of our results with those of Doebeli & Ispolatov (2014), we conform to their adaptive dynamics approach (also known as invasion analysis). Under this framework, the population is always fixed for a given phenotype, and evolution happens through a series of invasions of a population of resident phenotypes by better adapted mutants that differ by infinitesimally small phenotypic effects (Dieckmann & Law, 1996; Geritz *et al*., 1998). Since mutations are assumed to be rare, population dynamics are faster than evolutionary dynamics, such that the population size equilibrates at the carrying capacity of the resident before any new mutant invades (in other words, ecological and evolutionary time scales are decoupled).

Using the adaptive dynamics assumptions above, Doebeli & Ispolatov (2014) demonstrate that the invasion fitness *f*(**y**, **x**) of a mutant with phenotype **y** in a population fixed for the resident phenotype **x**, can be written as

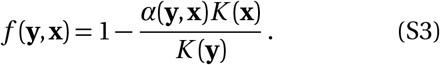

The rate and direction of evolutionary change at any time is then proportional to the selection gradient, i.e. the partial derivative of invasion fitness (*per capita* growth rate of a rare mutant) with respect to the invader’s phenotype (Dieckmann & Law, 1996; Geritz *et al*., 1998). If mutation effects are independent and of variance equal to one for all traits (which can be obtained by proper rescaling and change of coordinates, Martin & Lenormand, 2006; Doebeli & Ispolatov, 2014), the dynamics of evolution in the multidimensional phenotypic space depends solely on the multivariate selection gradient **s**(**x**) = (*s*_1_(**x**), *s*_2_(**x**),…, *s_d_* (**x**)). Each component *s_i_*(**x**) of this vector is the partial derivative of the invasion fitness in equation (S3) relative to trait *i* of the mutant, evaluated at the resident phenotype for that trait, which leads to

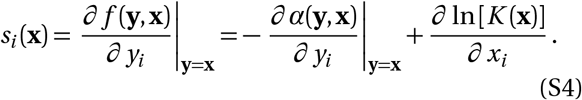

Following Doebeli & Ispolatov (2014), we assume that the interaction kernel in equation (S3) is such that its derivative in equation (S4) is a quadratic function of phenotypes. Note that this implies that *α*(**y**, **x**) itself (rather than its derivative with respect to y) is a third-order function of phenotypes, i.e. includes interactions between three traits, at least one of which belongs to the focal individual and one to its interactor. For simplicity, we do not consider interactions of even higher orders; including them would increase the likelihood of chaotic behavior due to the more complex feedbacks in the system (Ispolatov *et al*., 2015).

A final important point about this model is that, if the interaction kernel depended only on the phenotypic difference between interactors rather than on their actual phenotypes, that is, if we had α(**y**, **x**) = *f*(*z*) (where **z** = **y** – **x** and *f*() is a function that does not involve **x** nor **y**), then the corresponding term of the selection gradient in equation (S4) would be *f*’(0), which is not a function of **z** and thus cannot be a function of **x** and **y**. Hence the first two sums in equation (1) in the main text would be null if α(**y**, **x**) = *f* (**y**–**x**), which would preclude the occurrence of chaotic dynamics, since stabilizing selection alone does not produce chaos in such a model.

### The Lyapunov exponent

Chaotic dynamical systems can be identified by their Lyapunov exponents, which measure the rate of exponential increase in the distance between initially close trajectories in phenotype space (Ott, 2002). The main characteristic of chaotic systems is that, regardless of how close a set of trajectories start, trajectories initially diverge with time if they do not have the same exact initial state. Chaotic systems are, therefore, characterized by positive Lyapunov exponents (i.e. exponential rates of divergence). Dynamics that converge to cycles and fixed equilibrium states have zero and negative Lyapunov exponents, respectively. In multivariate systems, divergence can happen at different rates in different directions, but a positive rate of divergence in a single direction is sufficient for a system to be chaotic. Therefore, knowledge of the largest Lyapunov exponents is sufficient to identify chaos.

We numerically estimate the largest Lyapunov exponents of all our simulated trajectories as described by Sprott (2001). Given a previously simulated trajectory {**x**(0),**x**(dt),**x**(2d*t*),…}, we start by picking a point **z**_0_ positioned in a random direction at a distance *δ*_0_ = 10^−3^ from **x**(0). We then integrate the system from **z**_0_ for a single timestep d*t* the same way as done for the original trajectory. After this calculation, the distance *δ_f_* between the reached point **z**_*f*_ and **x**(dt) is recorded. We finally reset point **z**_0_ to lie in the same direction that separates **z**_*f*_ and **x**(dt), at a distance *δ*_0_ = 10^−3^ from **x**(dt), and proceed the integration from this point for another timestep. This process is iterated along the whole trajectory up to its end. The rate of divergence at each time step is calculated as

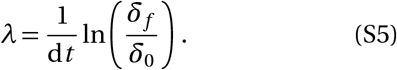

The estimate of the largest Lyapunov exponent is given by the asymptotic value of the average Lyapunov exponent as the number of considered time points tends to infinity, 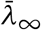. However, since we are interested in identifying transitions from chaotic to either periodic or equilibrium behavior, we used a local estimate of rates of divergence, 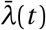, defined as the average *λ* in a window of 200 time units preceding time *t* (Supplementary Fig.S1). The length of this window was chosen such that there is enough smoothing of the short-term fluctuations of *λ*, while allowing for identification of transitions (as in Figure S1B). For simplicity, we refer to this measure in the main text as the Lyapunov exponent λ, omitting both the bar and *t*. Chaotic trajectories were easily distinguished from non-chaotic ones by visual inspection. Based on this criterion, we were able to categorize chaotic trajectories at a given time *t* if *λ*(*t*) was larger than a threshold *μ* = 0.05 under a constant optimum, and *μ* = 0.02 under an oscillating optimum. Trajectories were classified as converging to an equilibrium if 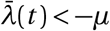, or to aperiodic cycle if 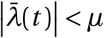.

**Fig. S1.**
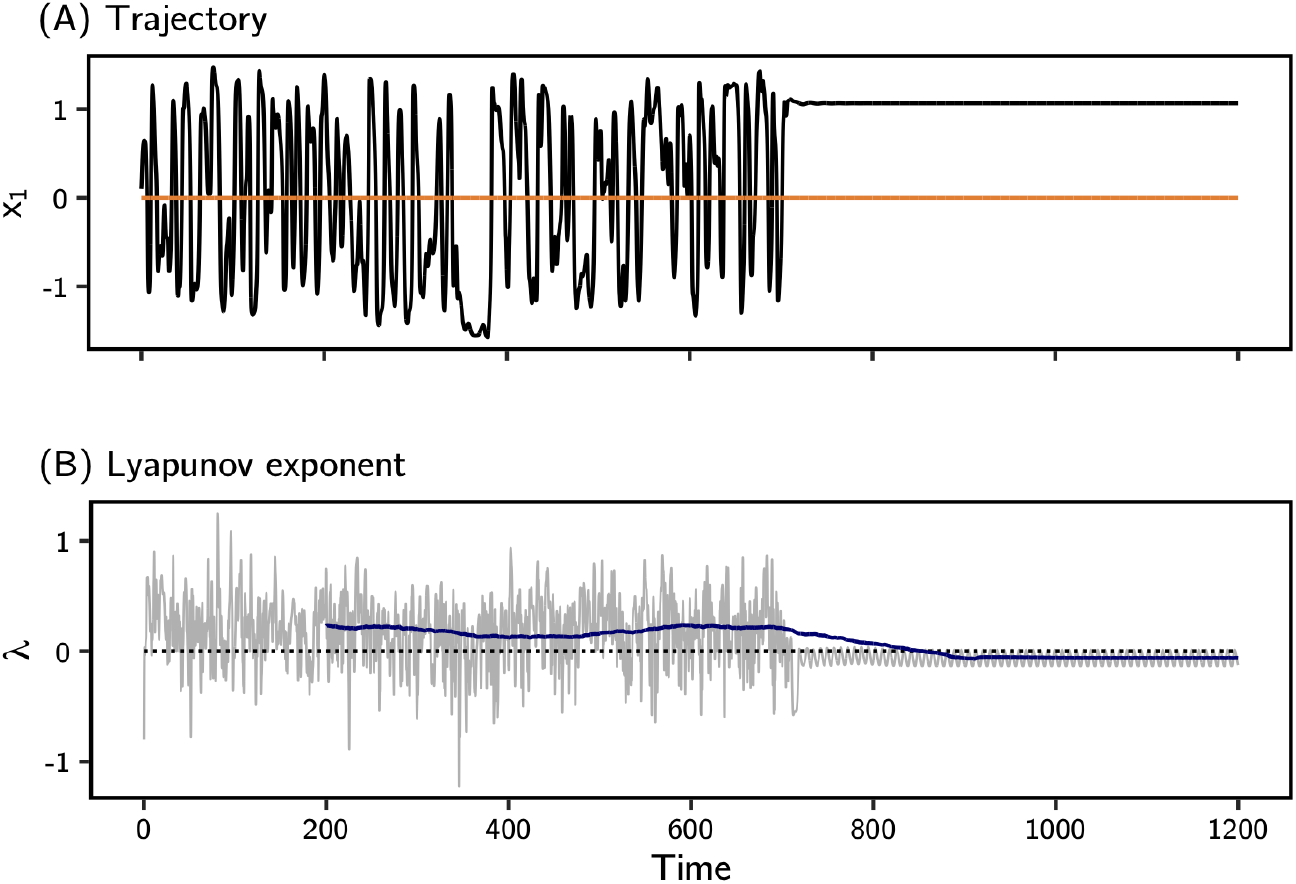
Calculation of the window-averaged Lyapunov exponent 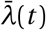. (A) The evolutionary trajectory for one single trait in a system of *d* = 70 is represented in black, under a constant optimum (orange). The dynamics transitions from chaos to a fixed point around time *t* = 700. (B) The Lyapunov exponent computed at each time point with equation (S5) varies substantially (gray line). However, when averaged over a window of 200 time units (in blue; calculated as described in the supporting text), the transition from chaos (*λ* > 0) to a fixed equilibrium (*λ* < 0) can be identified with some time lag.

**Fig. S2.**
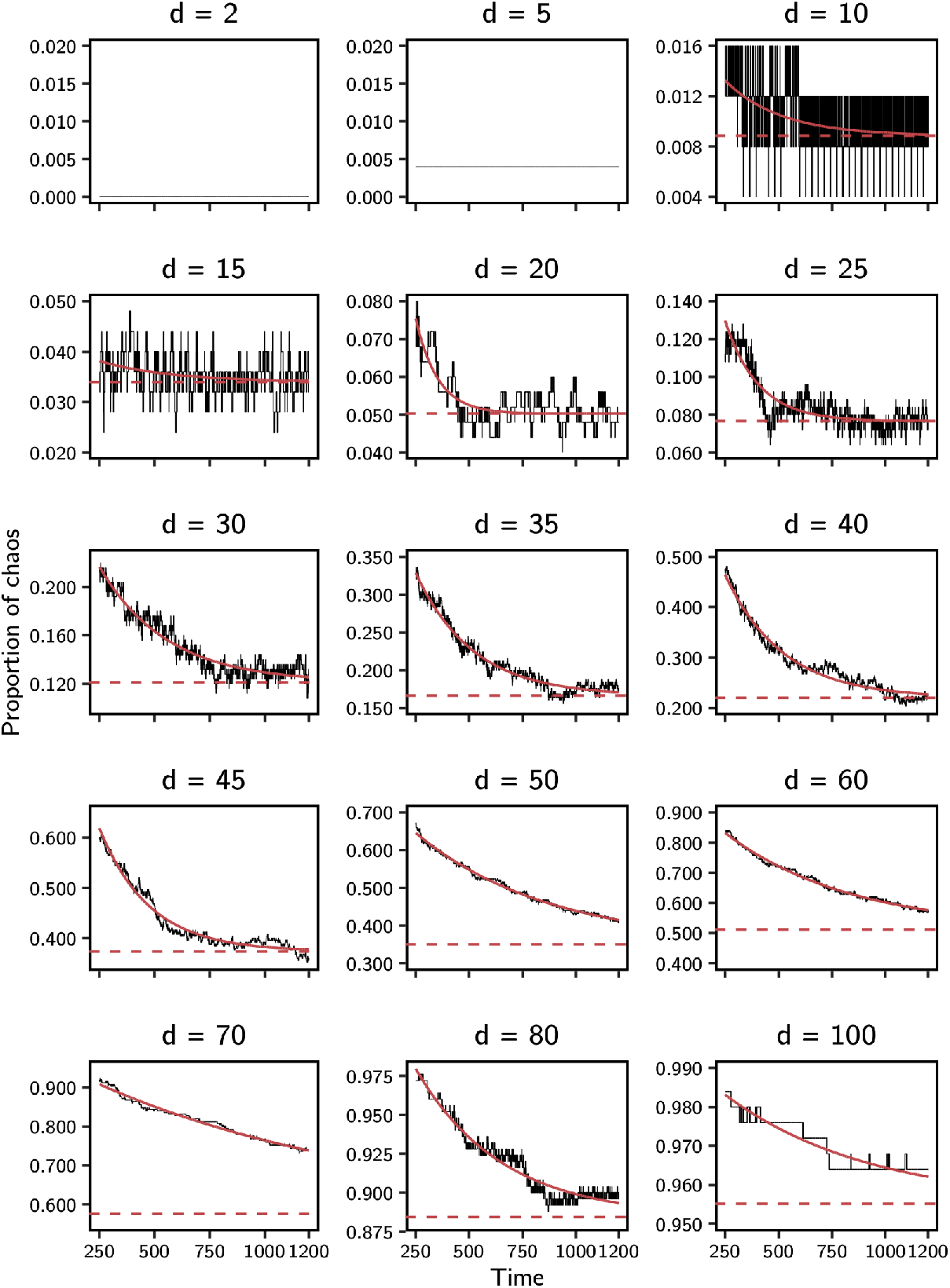
The proportion of trajectories categorized as chaotic (according to their window-averaged Lyapunov exponent *λ* (*t*)), out of 250 simulations under a constant optimum, decreases with time as some of them transition to either periodic cycles or fixed equilibria. Models of exponential decrease (red solid lines) were fit to the observed frequencies (as described in the Methods), and used to predict the asymptotic proportion of truly chaotic trajectories in infinite time (red dashed lines). Models were not fitted to dimensionalities *d* = 2 and 5 because of their insufficient number of chaotic trajectories. The black line in *d* = 5 is flat because the single chaotic trajectory observed did not transition during the simulation. The black lines in the other panels are jagged because the *λ* (*t*) of individual trajectories are only estimates that can switch between chaos and non-chaos in time.

**Fig. S3.**
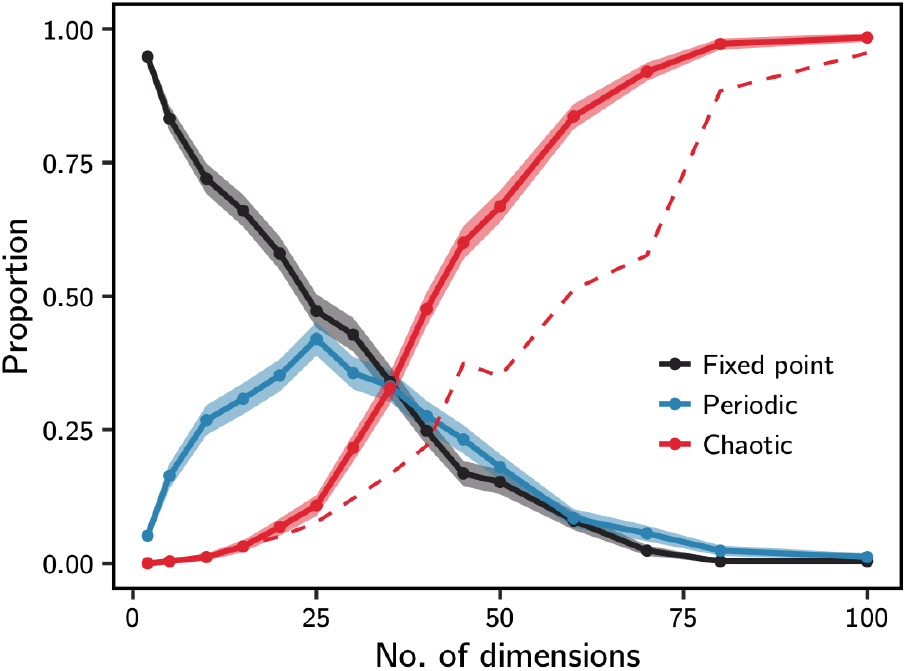
Organismal complexity and the probability of (transient) chaos in a constant environment. The proportion of trajectories identified as fixed point, periodic or chaotic at the beginning of the simulations (classified by their *λ*(250)) is represented in solid lines, with the standard errors of binomial proportions shown as shading. The proportion of trajectories identified as chaotic decreases with time, as transiently chaotic trajectories reach their eventual equilibrium fixed point or limit cycle. The red dashed line shows the asymptotic proportion of chaotic trajectories (at infinite time), as estimated from the exponential decrease, over *t* = 1200 time units, of the proportion of trajectories identified as chaotic (as described in the Methods and shown in Figure S2). Note that this curve is more noisy than the others, owing to error in estimation of the asymptotic values represented in Fig.S2. For each explored value of dimensionality *d*, estimations of frequencies of each type of dynamics were made based on 250 trajectories that were run up to *t* = 1200, as described in the main text.

**Fig. S4.**
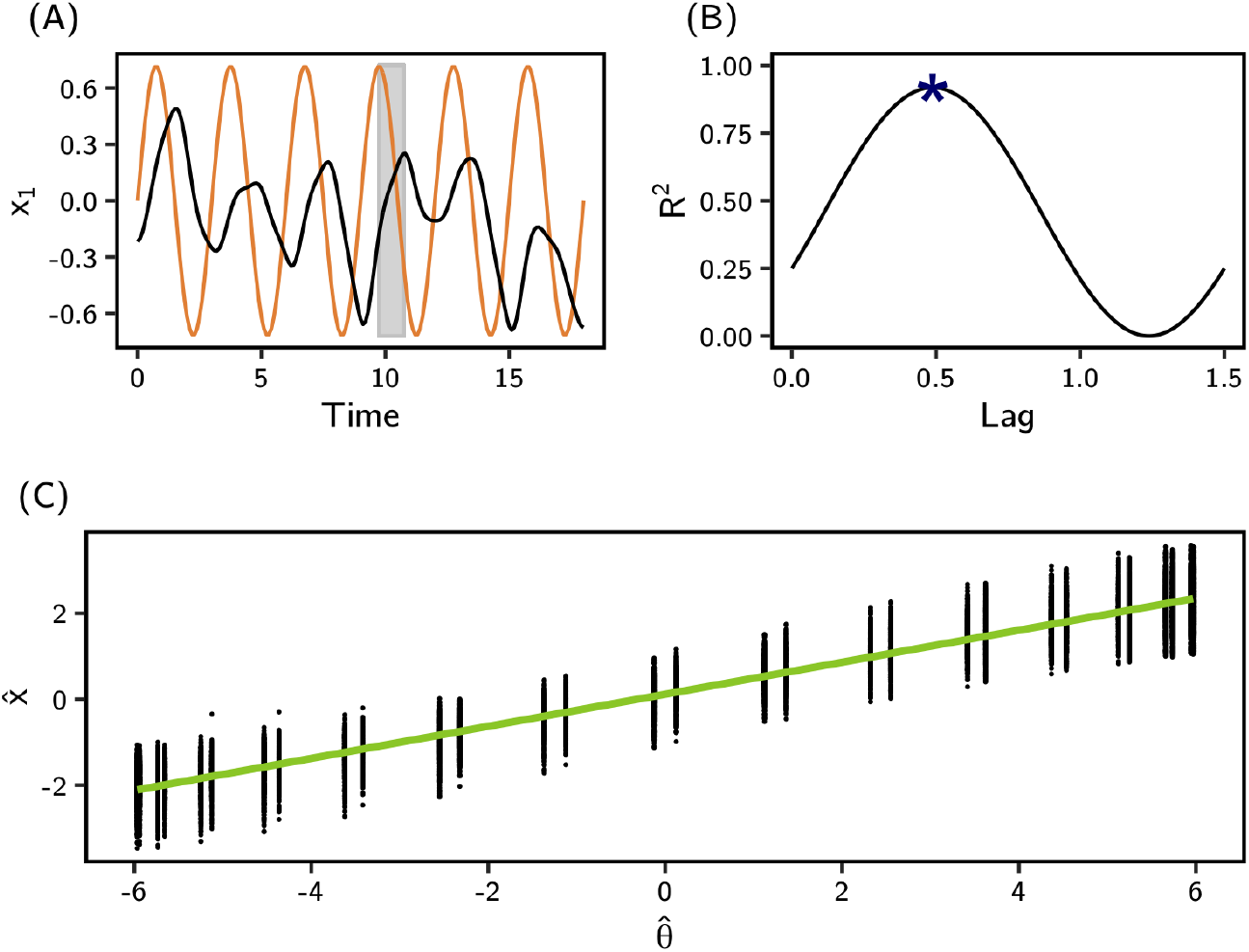
Correction of phenotypic lag in the regression of the phenotype on the optimum. (A) Even for a chaotic evolutionary trajectory (black line), part of the variation is caused by tracking of the optimal phenotype that moves in response to the oscillating optimum (orange line). This tracking happens with a certain temporal lag (represented by the shaded interval). (B) By regressing the phenotype 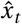 at time *t* on the optimum 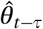 at an earlier time *t – τ* (both projected on the direction of environmental change in the optimum), the lag can be estimated by maximizing the *R*^2^ of the regression (marked with a star) with respect to *τ*, over half a period of optimum oscillation. (C) The relationship between 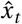 and 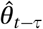 corrected by the lag is represented for simulated values (dots), together with the corresponding regression line shown in green. The slope of this line is expected to equal the ratio between the amplitude of the tracking component of phenotype and that of the oscillating optimum.

**Fig. S5.**
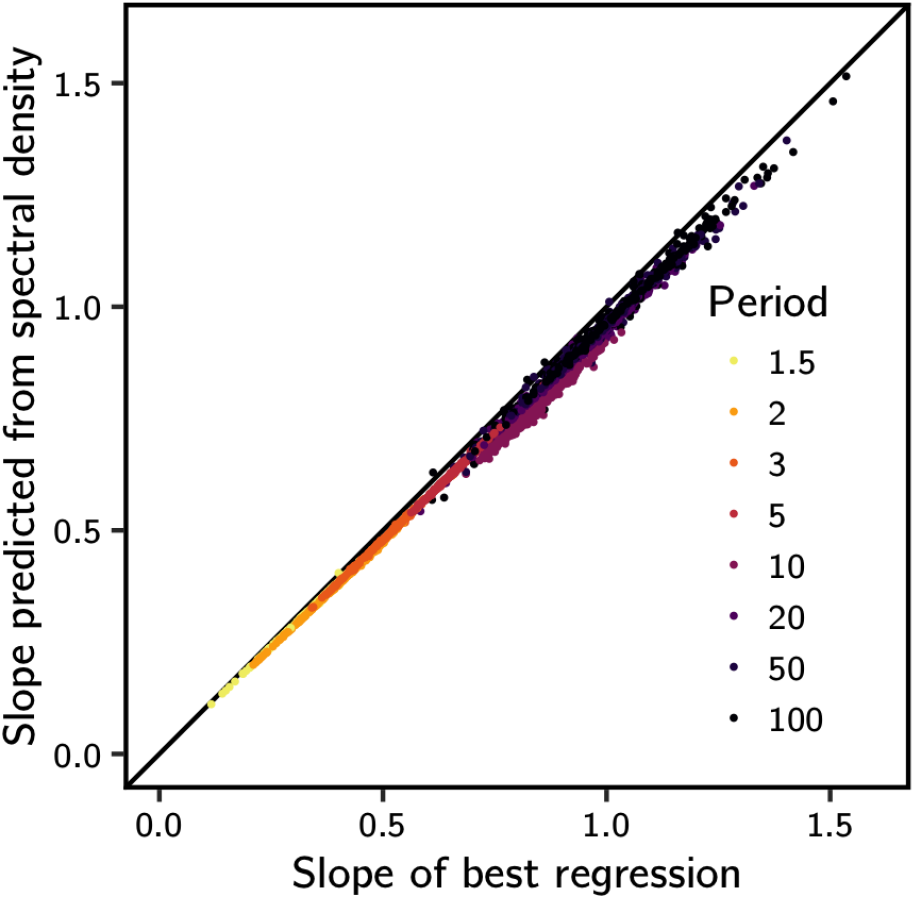
Estimating the regression slope from the spectral density. On the x-axis, the regression slope of the phenotype on the optimum (after correction for the lag, as explained in Figure S4) is expected to approximate the ratio between the amplitudes of the tracking component of the phenotype, identified by the linear regression model, and the amplitude of optimum oscillation (as shown in Figure S4C). In the spectral analysis of the phenotypic time series, the peak spectral density can be used to identify the period of the tracking component, which equals that of the oscillating phenotype (as shown in Figure 3C and D). Given that the spectral density at any period equals half the squared amplitude of the trajectory’s oscillation at that period, the amplitude of the tracking component of the trajectories can be estimated from this peak spectral density—as well as the ratio between this amplitude and that of the oscillating optimum. Here we show that this estimation, done for each trajectory independently (points, colored according to the period of optimum oscillation), closely approximates the slope of the regression models. However, this is only possible by interpolating the time-series of phenotype to increase the number of data points and, thus, improve the estimation of the spectral density at the period of optimum oscillation as done by the Fast-Fourier algorithm used here (as described in the Methods). We have made several of these estimations, each time linear-interpolating the time-series to a larger number of timepoints, one extra timepoint at a time up to the double of the original number of points. For each of these interpolated time-series, the peak spectral density was calculated, and the largest of these values (among all interpolated time-series of a single trajectory) was selected to estimate the amplitude of the component of phenotypic evolution that tracks the moving optimum.

**Fig. S6.**
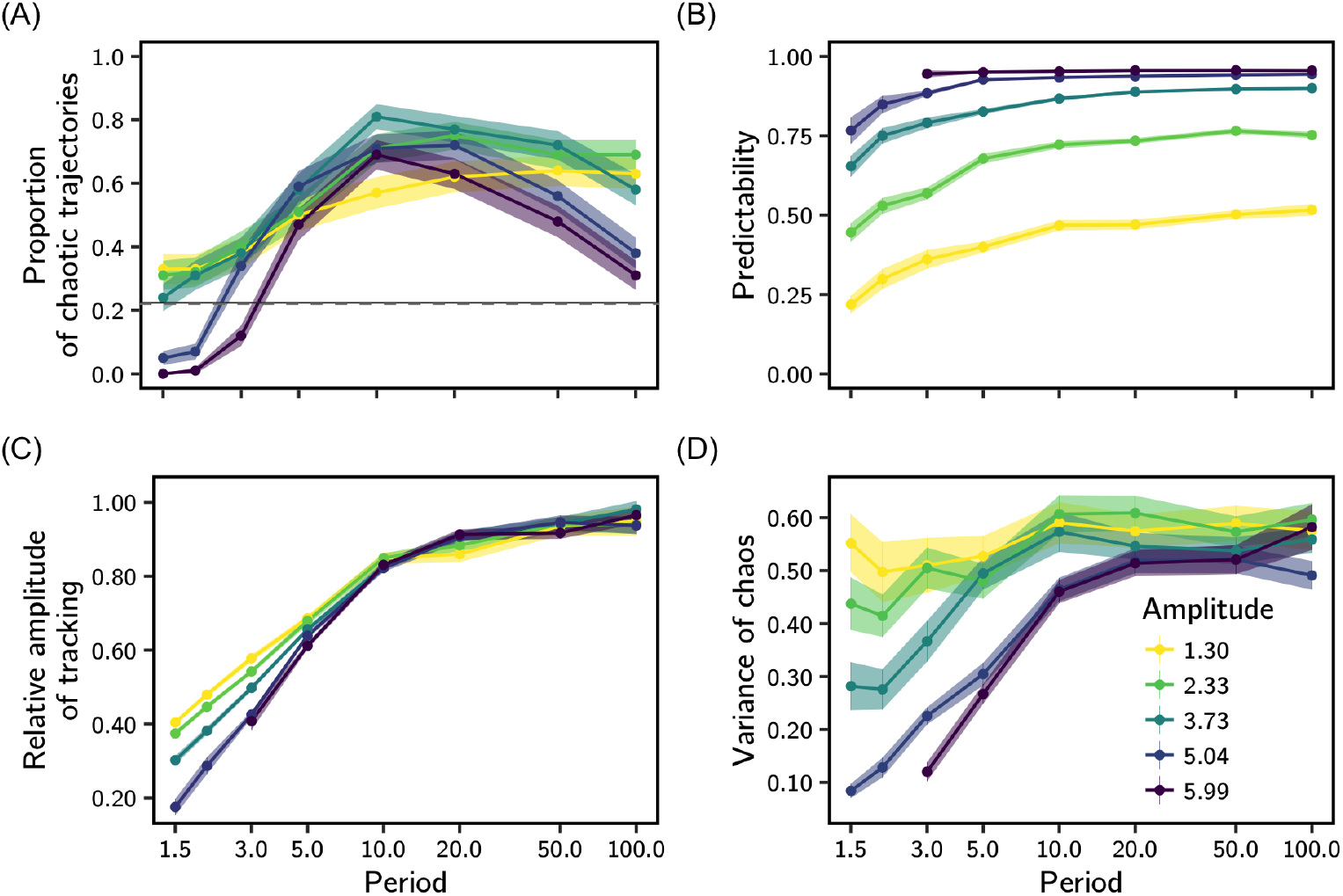
Predictability of evolution in a system of lower dimensionality. Here we replicate the oscillating optimum analyses presented in the main text, but for systems of *d* = 40. (A) Similarly to results for *d* = 70, the proportion of trajectories that are chaotic is decreased in relation to that of constant environment for conditions of short periods and large amplitudes of oscillation (complete loss of chaos even occurred for amplitude 5.99 and periods 1.5 and 2). This proportion increased for longer periods as for *d* = 70, but this increase was not monotonic (compare with Figure (2)). Results for (B) the predictability of chaotic dynamics, (C) relative amplitude of tracking, and (D) variance of chaos, are qualitatively similar to those for *d* = 70. The dashed lines in A have the same meaning as in Figure (2), and the variables represented in B-D are defined as in Figure 4. We show the average (lines and points) and standard error (shading) over 100 simulations for each combination of amplitude and period of optimum oscillation. The 100 sets of interaction coefficients and initial phenotypes used here were taken arbitrarily from the 250 sets used in constant environment simulations for the same dimensionality.

**Fig. S7.**
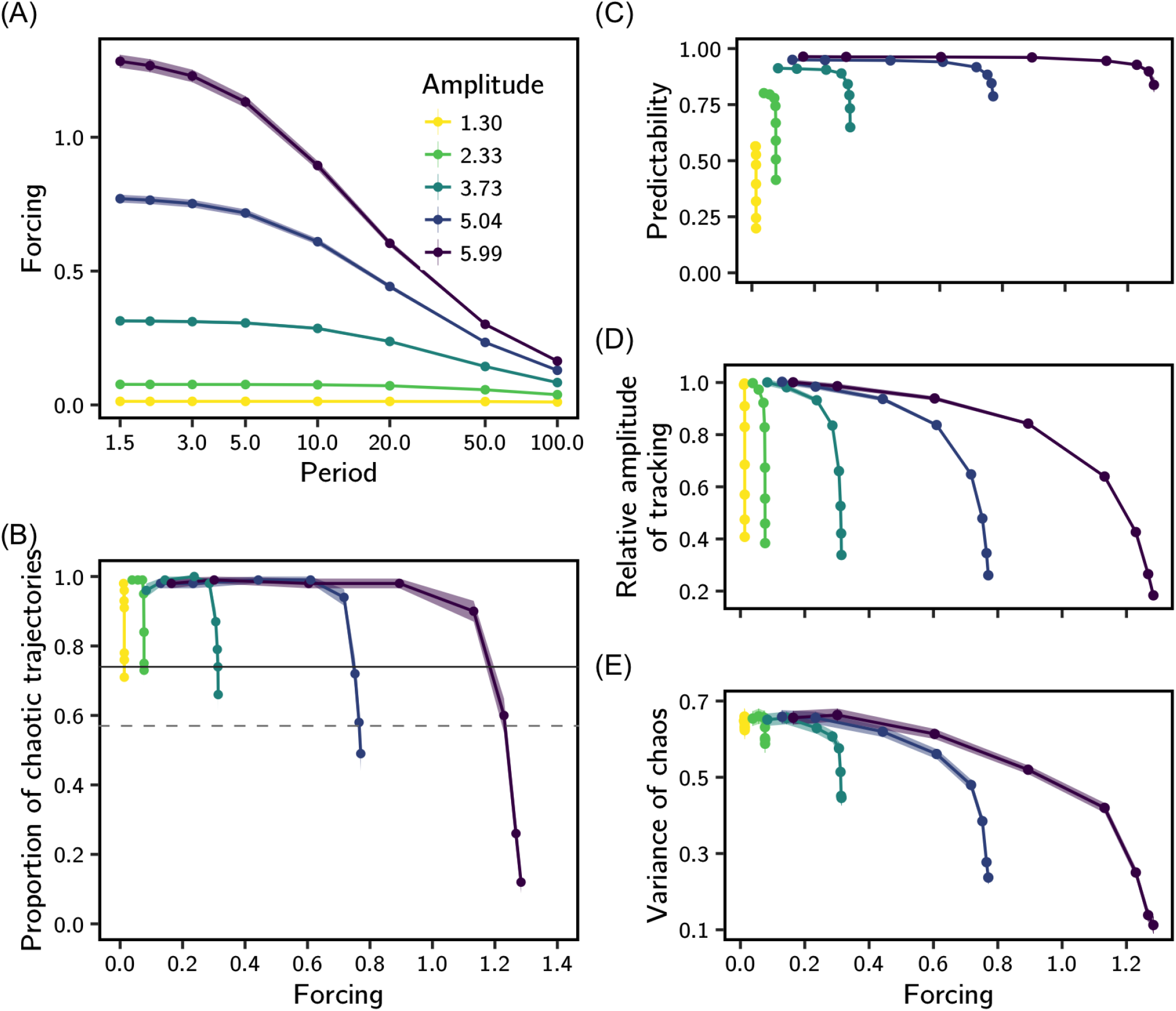
Effect of the strength of environmental forcing. We used a version of the model without frequency-dependent selection (interaction coefficients *b_ij_* and *a_ijk_* were set to zero) to measure how different patterns of optimum oscillation translate into intensities of environmental forcing on the system. Under frequency-independent selection, the evolutionary dynamics are led exclusively by the rightmost term in equation (S4). We defined forcing as 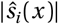, i.e. the absolute value of the scalar projection of the selection gradient in equation (S4) on the direction of environmental change (the diagonal of the phenotype space). This effectively measures the average magnitude of the selection pressure to which trajectories are submitted by the moving optimum. We solved the evolutionary dynamics numerically (as done in simulations with frequency dependence) until time *t* = 200, and computed the average forcing that trajectories experienced under the combinations of amplitudes and periods of optimum oscillation described in the main text. (A) A combination of short periods and large amplitudes of oscillation leads to the strongest forcing scenario. To assess whether this measure of forcing can partly explain our results *with* frequency-dependent selection, we plotted against it (B) the decrease in proportion of chaos (relative to a constant environment), (C) the predictability of evolutionary dynamics, (D) the relative amplitude of tracking, and (E) the variance of chaos. The dashed lines in B are the same as in Figure 2, and the variables represented in C-E are defined as in Figure 4. Each point represent the results for a single combination of amplitude and period of optimum oscillation. Points are colored by their respective amplitudes. We show the average (lines and points) and standard error (shading) over 100 simulations for each combination of amplitude and period of optimum oscillation.

**Fig. S8.**
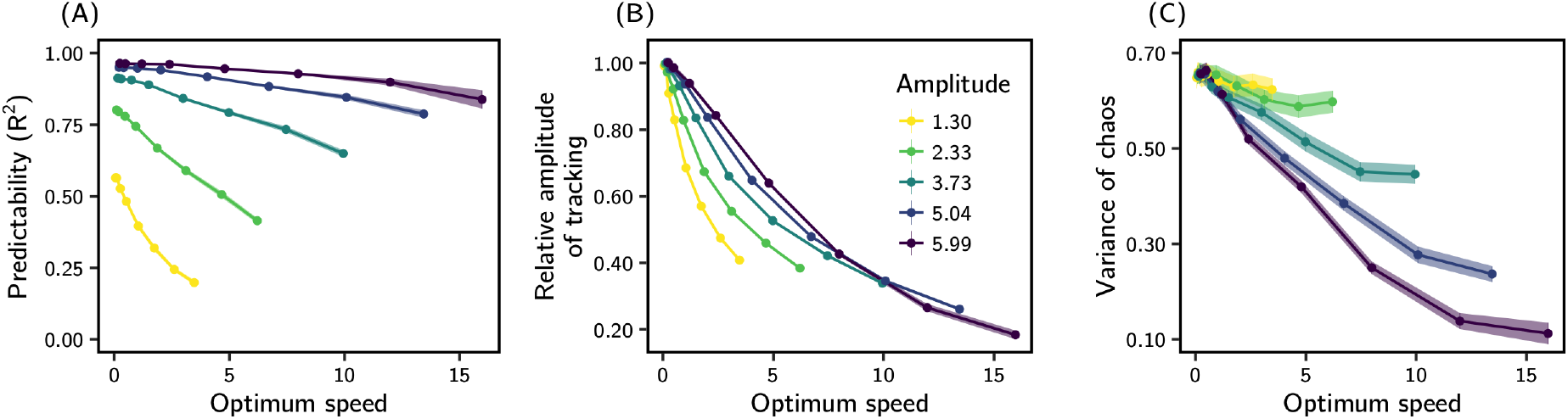
Linear regression of phenotype on the moving optimum against the average speed of optimum oscillation. The average speed of optimum oscillation (calculated from equation (2) as 4║**A**║/*T*) is higher for conditions of short period and high amplitude of oscillations. Predictability, relative amplitude of tracking and variance of chaos were measured as in Fig.4. We show the average (lines and points) and standard error (shading) over 100 simulations for each combination of amplitude and period of optimum oscillation.

**Fig. S9.**
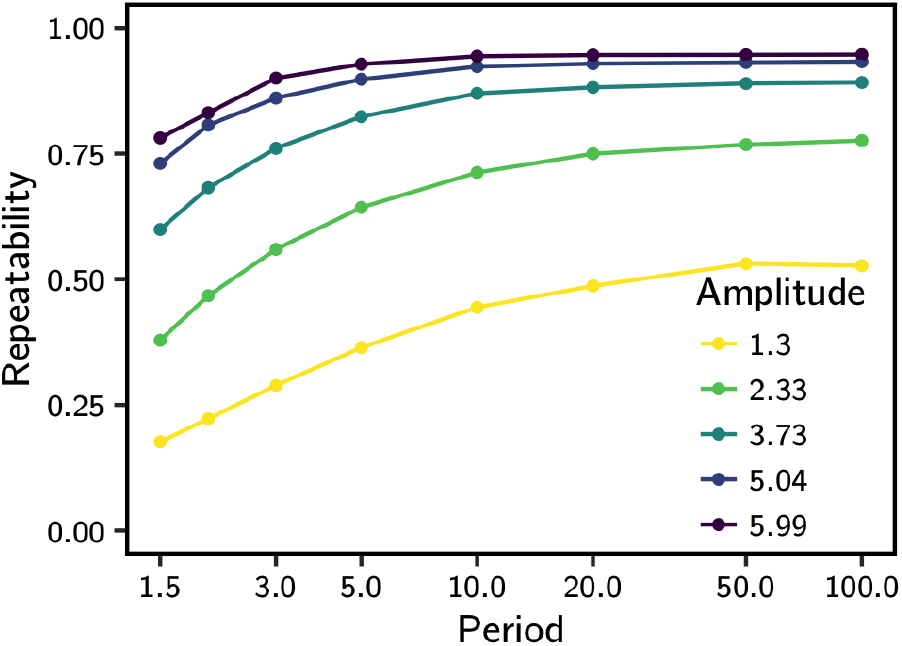
The repeatability of chaotic trajectories. The ratio of the variance through time of the mean trajectory (mean phenotype averaged across replicates) to the total variance of trajectories (across time and replicates) is shown for each condition of environmental oscillation. We used univariate phenotypic values obtained by scalar projections of the trajectories on the direction of optimum oscillation. The pattern observed is much similar to that of the predictability measured by the *R*^2^ of the linear regression of the phenotype on the optimum (Fig.4 in the main text). Repeatability differs from our measure of predictability because it does not rely on knowledge of the optimal phenotype.

**Fig. S10.**
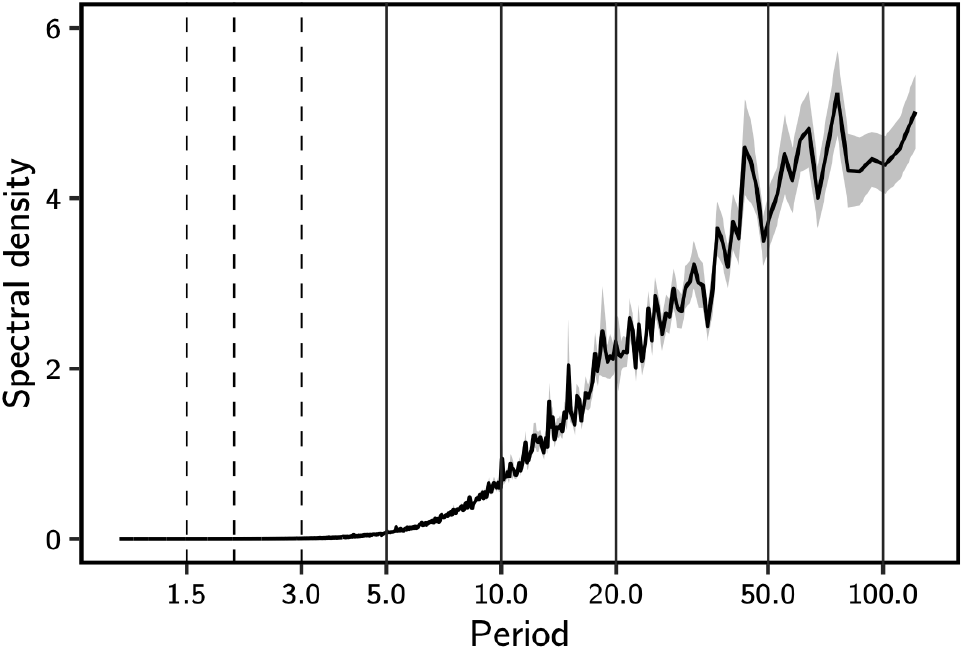
The average spectral density of evolutionary trajectories in constant environment describes the typical range of periods at which the phenotype oscillates. Among all explored values of period of optimum oscillation (vertical lines) those that led to reduction in the proportion of chaos in relation to the constant environment (as seen in Figure 2, and here marked by vertical dashed lines) are much shorter than the typical oscillations of the system in constant environment. The mean spectral density (solid wiggly line) and its associated standard error of the mean (shading) were calculated based on the 250 trajectories run in constant environment for *d* = 70. The spectrum was calculated as described in the main text, for the projection of the phenotypic vector along the diagonal of the phenotype space.

